# Learning to Move and Plan like the Knight: Sequential Decision Making with a Novel Motor Mapping

**DOI:** 10.1101/2024.08.29.610359

**Authors:** Carlos A. Velázquez-Vargas, Jordan A. Taylor

## Abstract

Many skills that humans acquire throughout their lives, such as playing video games or sports, require substantial motor learning and multi-step planning. While both processes are typically studied separately, they are likely to interact during the acquisition of complex motor skills. In this work, we studied this interaction by assessing human performance in a sequential decision-making task that requires the learning of a non-trivial motor mapping. Participants were tasked to move a cursor from start to target locations in a grid world, using a standard keyboard. Notably, the specific keys were arbitrarily mapped to a movement rule resembling the Knight chess piece. In Experiment 1, we showed the learning of this mapping in the absence of planning, led to significant improvements in the task when presented with sequential decisions at a later stage. Computational modeling analysis revealed that such improvements resulted from an increased learning rate about the state transitions of the motor mapping, which also resulted in more flexible planning from trial to trial (less perseveration or habitual responses). In Experiment 2, we showed that incorporating mapping learning into the planning process, allows us to capture (1) differential task improvements for distinct planning horizons and (2) overall lower performance for longer horizons. Additionally, model analysis suggested that participants may limit their search to three steps ahead. We hypothesize that this limitation in planning horizon arises from capacity constraints in working memory, and may be the reason complex skills are often broken down into individual subroutines or components during learning.

## 1. Introduction

During their lifetime, humans can develop a wide and complex repertoire of motor skills such as dancing, swimming, riding a bicycle, playing musical instruments, or playing video games. The intricate nature of these activities has made their scientific study equally complex. First, humans must figure out the motor commands that lead to the desired outcomes. For example, which configuration of the hand produces a given chord on the guitar or what button presses make a character jump or walk in a video game? The learning of these mappings poses the first challenge when attempting to acquire a novel motor skill (Fitts & Posner, 1967; Adams, 1971; Ackerman, 1988; Newell, 1985, 1991). A second challenge arises as the majority of complex skills are extended in time, involving sequencing together a set of actions with the mapping to accomplish goals. For example, a given combination of chords is necessary to generate a song, and sequences of button presses make players navigate through different levels of a video game. The dependence of goals on the concatenation of actions gives rise to one of the key processes of human cognition: planning (Hunt et al., 2021; Mattar and Lengyel, 2022). In this study, we aim to understand how complex motor skills — requiring both sequential decision-making and the learning of a novel mapping — are acquired.

Developments on motor learning and planning research currently provide no clear answers to this problem. On the one hand, the acquisition of a novel mapping has been studied in sequence learning tasks where people have to learn what action to take, normally key presses, when arbitrary visual stimuli are presented on a computer screen (Balsters & Ramnani, 2011). However, there is generally no overarching goal towards which people can freely use the mapping, i.e., choosing their own sequence of actions, such as in video games. Additionally, in most sequence learning tasks, the sequence to be learned is fully specified by the experimenter (Korman et al., 2003; Kami et al., 1995). This limits considerably the planning aspect of the tasks, especially for internally-generated skills such as in musical improvisation or playing video games (Bera et al., 2021a, 2021b)

One exception is the work on grid-world navigation (Fermin, et al., 2010, 2016; Bera et al., 2021a, 2021b; Franklin and Frank, 2018, 2022; Velázquez-Vargas et al., 2024), where participants have to move a cursor from start to target locations using keys with an arbitrary movement-mapping. In these tasks, the learned mapping can be freely manipulated to generate sequences of actions to arrive at the targets, closely resembling video game playing. However, an important limitation of these studies is the small set of stimuli (start and target locations) that is commonly used, which makes it likely that participants readily use memorized solutions rather than performing planning (Velázquez-Vargas et al., 2024). Additionally, given that the mappings are typically simple, learning proceeds rather quickly, reaching ceiling performance within a few trials (Fermin et al., 2010, Bera et al., 2021a; Velázquez-Vargas et al., 2024). This contrasts with the complexity of the motor skills that humans can acquire, where the mapping can take hours or years to be fully learned and where planning processes are constantly being deployed due to the changing goals.

On the other hand, planning research, which focuses on how biological and artificial agents maximize their future rewards, has progressed significantly through the study of multi-step tasks (Solway and Botvinick, 2015; Gläscher et al., 2010; Daw et al., 2011) and navigation tasks (Tolman, 1948; Redish, 2016; Mattar and Daw, 2018; Johnson and Redish, 2007). In humans, various algorithms inspired by artificial intelligence (AI) have been proposed as hypotheses for how planning occurs (van Opheusden et al., 2017, 2023; Élteto et al., 2023; Krusche et al., 2018). While the specific algorithms humans use remain unknown, it is likely that the brain, given its limited resources, implements heuristics to reduce computational costs of planning, such as pruning or truncating the search of future scenarios (for a review, see Mattar and Lengyel, 2022).

A key component of most planning algorithms is the existence of an internal model of the world, which evaluates candidate actions based on knowledge about the rewards and state transitions. This model can be known beforehand such as in board games (van Opheusden et al., 2021) or learned through experience as in instrumental learning experiments (Glächer et al., 2010; Daw et al., 2011). In most planning tasks, however, the internal model is assumed to require minimal or no motor learning at all. For instance, participants in multi-step choice tasks may only need to click on the intended choice to transition to a different state. Similarly, animals tested in mazes rely on a well-calibrated mapping of their body movements to navigate and reach rewards. While this assumption is reasonable for various cognitive tasks such as playing board games or performing spatial navigation, it contrasts with how motor skills are developed, where a mapping between state transitions and motor commands must also be learned.

In the present work, we developed a grid navigation task to study planning when state transitions require motor learning, while addressing the aforementioned limitations. In our experiments, participants moved a cursor from start to target locations using eight keyboard keys. Each key moved the cursor in L-shaped directions, following the movement of the chess piece known as the “Knight” (Velázquez-Vargas and Taylor, 2023). This mapping could be more closely comparable to the complexity of mappings humans acquire in real world scenarios, involving multiple actions with a non-trivial rule.

In Experiment 1, we show that learning the motor mapping in the absence of planning leads to performance benefits on subsequent stages of the task, where sequential decision making is needed. In addition, in Experiment 2, we show that incorporating mapping-learning into the planning process allows us to explain differences in performance for distinct planning horizons.

## 2. Methods

### 2.1. General

#### 2.1.1. Participants

Seventy five undergraduate students (34 males, 39 females and 2 non-binary; mean age = 19.6, sd = 1.5) from Princeton University were recruited through the Psychology Subject Pool. The experiments were approved by the Institutional Review Board (IRB). All participants provided written informed consent before performing the experiment.

#### 2.1.2. Apparatus and task design

The experiments were carried out in person using the same computer equipment for all participants. Stimuli were displayed on a 60 Hz Dell monitor and computed by a Dell OptiPlex 7050’a machine (Dell, Round Rock, Texas) running Windows 10 (Microsoft Co., Redmond, Washington). Participants made their responses using a standard desktop keyboard. All experiments were run on the browser and hosted on Google Firebase. Subjects were seated in front of the computer and were asked to follow the instructions to begin the task.

In a 15×15 grid, participants were asked to move a cursor in the form of a frog from start to target locations represented by clovers (Figure 1A). The cursor moved deterministically based on the movement rule of the piece of chess known as the Knight. The Knight moves in L-shape directions: either two squares in one direction (up, down, right, left) followed by one square perpendicular to that direction, or one square in one direction followed by two squares perpendicular to that direction. Participants were not informed about which key moved the cursor in a particular direction. On a given position of the grid, the locations where the cursor could move to were highlighted in color in order aid learning. From a pilot study we found out that without this shading participants’ performance was significantly worse, although they could learn eventually. Subjects moved the cursor using eight keys of their keyboard (A,S,D,F,H,J,K,L), which were randomly associated with a unique Knight move at the beginning of the experiment.

**Figure 1.**
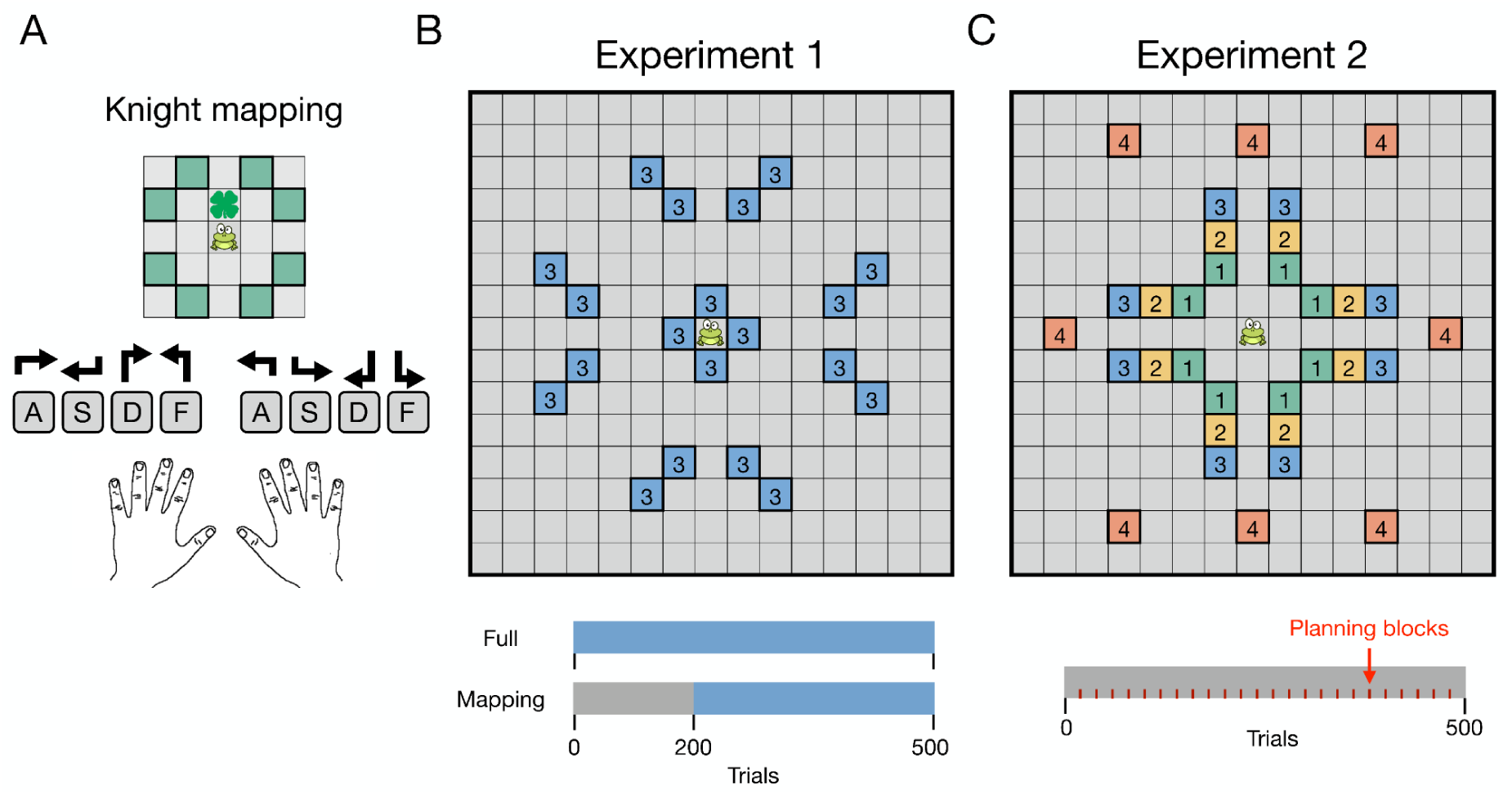
Grid navigation with the Knight mapping. **(A)** Participants were instructed to move a frog from start to target locations using eight keys, each corresponding to one of the Knight’s possible moves in chess. The Knight moves in an L-shape directions: either two squares in one direction (up, down, right, left) followed by one square perpendicular to that direction, or one square in one direction followed by two squares perpendicular to that direction. To facilitate learning, the possible states the frog could move to at any given time were highlighted throughout the experiment. A single target location, out of 20 possible, was presented on every trial in the form of a clover **(B)** In Experiment 1, participants were grouped in two groups. In the Full group, they performed 500 trials with the targets being three moves away. Participants did not see the number indicating the distance to the target. In the Mapping group, participants experienced 200 trials with target locations being one move away in order to encourage the learning of the mapping. Subsequently, they performed 300 trials with the targets placed three moves away as in the Full group. **(C)** In Experiment 2, target locations were placed either one, two, three or four moves away from the start location. Every 20 trials, participants experienced a “planning block” consisting of four trials (one for each target distance) where no movement feedback was provided. During these trials, the frog turned blue, and participants were informed that it had frozen and it would not be able to move, but that they should still attempt to arrive at the target.

On every trial, the cursor appeared at the center of the grid and a target location was indicated with a clover. The distance to the target varied by experiment (see below). Importantly, only one pair of start-target locations appeared on a given trial. If participants arrived at the target using the minimum number of moves, they received one hundred points. Then, they would lose five points for every extra move. If they did not arrive at the target location in ten seconds, it was considered a miss, they received zero points and moved on to the next trial. This points system has been used in previous grid navigation studies (Bera et al., 2021a). At the end of each trial participants observed the points obtained as well as the cumulative points since the beginning of the experiment.

### 2.2. Experiments

#### 2.2.1. Experiment 1

The goal of this experiment was to test whether the acquisition of complex motor skills, involving sequential decision making, can be benefited from learning of the motor mapping first, followed by the incorporation of planning at a later stage. In order to tackle this question, participants performed the grid navigation experiment previously described in two groups. In the Full group (n = 25), participants performed 500 trials and the target locations appeared always three steps away from the starting point (Figure 1B). We hypothesized that participants in this group face a double challenge: they need to arrive at target locations that require planning, and they need to do so while learning a non-trivial mapping. In order to reduce the likelihood that participants memorized the sequences to the targets and encourage planning, we used twenty different targets. They were presented in a random order, and each of them had to appear before showing them up again. The latter manipulation prevented the same target from being repeated multiple times. In the Mapping group (n = 25), participants first experienced 200 trials with target locations being one move away in order to encourage the learning of the mapping.

Participants were not informed that the first 200 trials had this purpose. Crucially, for this initial period, the trial was always terminated after the first move, such that participants could not generate further key presses to attempt to arrive at the target. Therefore, they could generate a maximum of 200 moves in the first 200 trials. Subsequently, they performed 300 trials with the targets placed three moves away as in the Full group.

#### 2.2.2. Experiment 2

The goal of this experiment (n = 25) was to directly assess planning processes throughout the course of learning a complex motor skill. In particular, we asked whether key signatures of planning such as differences in performance with varying distance to the targets (Jensen et al., 2024), would manifest in our task. In order to do so, we incorporated two main modifications. First, we manipulated the planning horizon by presenting targets that were either one to four steps away from the starting point. Participants experienced one trial from each horizon at randomized order before experiencing a target with the same horizon again. The target locations were randomly chosen out of 32 unique locations (eight for each horizon; Figure 1C). In addition, every 20 trials we incorporated a “planning block” where no movement (state transitions) or performance (points) feedback was provided. In particular, participants were instructed that in some trials the frog would turn blue, indicating that it froze and would be unable to move. However, they should still make the responses attempting to reach the target location. These blocks consisted of four trials, one for each planning horizon, and were presented at a random order. The goal of the planning blocks was to encourage planning prior to generating the sequence as online corrections were not possible or significantly harder to make, i.e., based on the believed location of the frog. We hypothesized that this manipulation would maximize the effects generated by the different planning horizons. Importantly, unlike in the first 200 trials in the Mapping group of Experiment 1, the trials in Experiment 2 could only be terminated by arriving at the target or by exceeding the time limit (ten seconds). This was also true for targets that were one step from the starting point, which allowed participants to continue attempting to arrive at the target even if they missed in the first move.

### 2.3. Computational models

To provide further insight into the cognitive processes involved in acquiring complex motor skills requiring sequential decision making, we evaluated a set of computational models based on three components which have provided convincing evidence about their influence on behavior: planning (Hunt et al., 2021; Mattar and Lengyel, 2022; Tolman, 1948), reward learning (Sutton and Barto, 1998; Schultz et al., 1997), and action perseveration (Miller et al., 2019, 2021; Daw et al., 2011; Thorndike, 1911).

#### 2.3.1. Model-based planning (MB)

Due to the sequential nature of our task, we hypothesized that participants could estimate the value of their actions based on simulations about their future outcomes. In order to do so, we implemented a constrained version of A* (“A-star”; Hart et al., 1968) as a planning model. A* is a widely used search algorithm with various applications such as in GPS navigation and in video-games (Rios and Chaimowicz, 2010). It is particularly well-suited for deterministic tasks like ours, offering significant computational advantages over uninformed algorithms such as Breadth-First Search (BFS; Moore, 1959) and Depth-First Search (DFS; Nilsson, 2014). These advantages come from A*’s ability to efficiently balance path cost and heuristic information, leading to faster and more optimal solutions. Specifically, in A* the cost *f* of a state *s* is defined by:

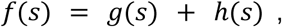

where *g*(*s*) is the cost (number of steps) from the start state to the *s* state. *h*(*s*) is a heuristic function, which provides an estimate of the cost from state *s* to the target. In A* search, paths are explored based on a priority queue where states with lower cost are prioritized.

As a heuristic function *h*(*s*), we used the chessboard distance:

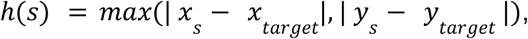

where *x* and *y* represent the coordinates of the state *s* and the target. The chessboard distance provides an intuitive variable to guide planning in our task, as it simply involves counting the number of grid cells away from the target. Subsequently, we can calculate the cost *C* of transitioning to state *s_i_* (where *s_i_* represents any of the neighboring states to which the cursor can move to in the next step, following the Knight rule):

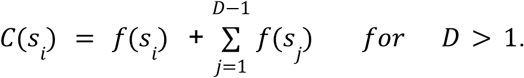

Here, the first term represents the immediate cost of moving to *s_i_*, while the second term represents the cumulative cost of the path starting from *s_i_* as determined by A* search. *D* is the depth of search, and is meant to capture the finite number of steps that humans plan into the future. In a model that considers only the next step *C*(*s_i_*) = *f*(*s_i_*). For Experiment 1, we set *D* = 3, which corresponds to the initial distance to the target, as in this experiment we were mostly interested in the learning of the transition probabilities and previous work has shown that this value falls within the range of the number of steps humans plan into the future (Huys et al., 2015; van Opheusden, 2023; Arad and Rubinstein, 2012; Snider et al., 2015). However, in Experiment 2, where we evaluated planning directly by manipulating the planning horizon, we compared models with different values of D.

Notably *C*(*s_i_*) only provides the cost of moving to the available states, but not how to do it. The mapping between the keys and the state transitions still needs to be learned. In previous work (Velázquez-Vargas et al., 2024), we proposed that this learning process can be modeled using Bayes’ rule. In this approach, each state transition is treated as a category, with new observations updating the beliefs about the true transitions through a Dirichlet-categorical model. However, an important limitation of this approach is that it does not account for differences in the learning rates of the state transitions, whether by individuals or due to experimental conditions. To address this, we adopted an alternative update rule based on prior work on reinforcement learning (Lee et al., 2014). Here, the transition probability from state *s* to state *s_i_* after taking action *a_k_* is given by:

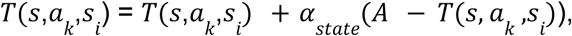

where α_*state*_ is the learning rate for the state transitions. A = 1 if action *a_k_* is chosen and A = 0, otherwise. Importantly, in our task *T*(*s,a_k_, s_i_*) is the same for all *s_i_*, as the mapping does not change depending on the location on the grid. *T*(*s*,*a_k_*, *s_i_*) was initialized as 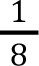, indicating that the eight transitions from the Knight rule were equally likely for all keys at the beginning of the experiment. Subsequently, the cost of acton *a_k_* is computed as:

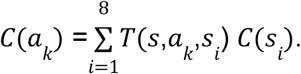

Finally, the probability ϕ of selecting action *a_k_* is generated using a Softmax function and − *C*(*a_k_*), so actions with less cost are more valuable:

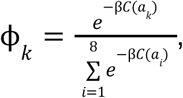

where β is the inverse temperature parameter. This model has two free parameters: α_*state*_ and β.

#### 2.3.2. Model-free learning (MF)

This model assumes that participants update the value of their actions using reward prediction errors (Sutton and Barto, 1998; Schultz et al., 1997; Rescorla and Wagner, 1972; Niv and Schoenbaum, 2008). In particular, we implemented SARSA (Sutton and Barto, 1998), a common temporal difference model:

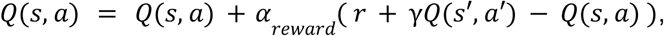

where *Q*(*s, a*) is the current value of the state-action pair, α_*reward*_ is the learning rate and *r* the rewards, which corresponded to the actual points at the end of the experiment. γ is a discount factor and *Q*(*s*′, *a*′) the estimated value of the next state-action pair. Importantly, in order to reduce the computational demand of storing state-action pairs for each target location, we used a state representation based on relative coordinates with respect to the target. This way, the state-action values can be reused across different targets. Lastly, the probability ϕ of selecting an action is generated using a Softmax function:

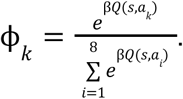

This model has three free parameters: α_*reward*_, γ and β.

#### 2.3. 3. Action perseveration (Persev)

Previous studies have shown that actions can be strengthened merely through repetition, independent of the reward obtained (Miller et al., 2019, 2021; Daw et al., 2011; Thorndike, 1911), leading to the formation of habits. Given the relatively infrequent rewards compared to the number of key presses in our task, we hypothesized that the perseveration on previous actions could be a contributing factor on participants’ choices. In particular, we implemented an update rule based on Miller et al., (2021):

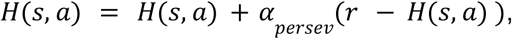

where *H*(*s,α*) is the perseveration value of the state-action pair, α_*persev*_ has a similar function of a learning rate, and determines the influence of past choices, with values closer to 1 indicating a greater weight of the most recent choices. Here, r = 1 if the action for the given state-action pair is selected, and r = 0, otherwise. Similar to the MF algorithm, we adopted a state representation based on relative coordinates. The probability ϕ of selecting an action is then generated using a Softmax function:

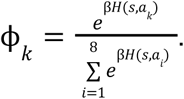

This model has two free parameters: α_*persev*_ and β.

#### 2.3.4. Hybrid models

We considered the possibility that each of the above-mentioned components could influence participants’ behavior. Specifically, we propose that responses could be the result of a linear combination of the MB, MF and perseverative components. We evaluated three hybrid models with two components and the full model with the three components. In the three-component hybrid, the value *U* of actions on a given state is given by:

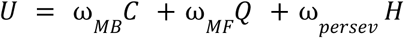

where ω_*MB*_, ω_*MF*_ ω_*persev*_ are free parameters with values between [-1,1]. In order to have values of C, Q and H that were within the same range, we normalized them to be between [0,1]. For the two-component hybrid models, one of the three terms was omitted. The probability ϕ of selecting an action is generated using a Softmax function:

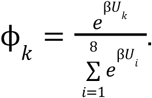

The number of free parameters in the hybrid models ranges between 5-8 depending on the components incorporated.

### 2.4. Data analyses

#### 2.4.1. Behavioral results

In Experiment 1 and 2, we used the points obtained in the task over trials as a measure of performance. In addition, in Experiment 1, in order to control for the difference in the number of key presses between the groups, we also computed the points by key presses. To do so, we grouped the key presses in bins of 40, and for each subject we obtained the averaged points they obtained in that period. Importantly, only the last responses in a trial can earn points. For example, if in 40 key presses a subject arrived at the target 10 times, the score for the bin corresponded to the average of the points obtained in those 10 times.

#### 2.4.2. Model analyses

The parameters from our models were estimated at the individual level using Bayesian Adaptive Search (Acerbi and Ma, 2017) implemented in Matlab code. We verified that the model parameters of the best fitting model (MB-Persev) were recovered reliably (Figure S1), and that the tested models with different depths were identifiable among each other (Figure S2). For Experiment 2, model fitting was based on the trials with feedback as in the planning blocks the states were no longer observable, and it is unknown where the updates from learning would be attributed to. Simulations on Figure 6 were performed by randomly sampling 100 target locations from the ones that participants could encounter. In addition, we used the mean of the best fitting parameters from the Full group in Experiment 1, with the exception of the inverse temperature parameter which we set to β = 15, to reduce exploration. The values estimated for β from the Full group were near 1, however, the model was fit on responses while we were trying to predict the outcome of the entire sequence (trial points). Simulations on Figure 7 were generated in the same way but using the best fitting parameters in Experiment 2.

## 3. Results

### 3.1. Experiment 1

Acquiring new motor skills can be challenging since they often require simultaneous learning of the motor mapping and planning. In this experiment, we asked whether separating the processes of learning and planning can speed skill acquisition. To do so, two groups participants performed a grid navigation task where the goal was to move a cursor from start to target locations using eight keyboard keys, which were mapped to one of eight directions according to the Knight rule in chess (Figure 1A). If participants arrived at the target using the minimum number of key presses, they received 100 points and lost 5 points for every extra move. In the Full group (n = 25), participants performed the experiment while attempting to arrive at target locations that were always three steps away. We hypothesized that in this group, participants faced a double challenge: learning a complex mapping and performing sequential decision making. In contrast, the Mapping group (n = 25), first experienced 200 trials with target locations placed one step away to encourage the learning of the mapping before having to plan. Importantly, during this period the trials were terminated after the first move such that participants could generate a maximum of 200 moves. Subsequently, participants performed 300 trials with the target locations being three steps away as in the Full group.

#### 3.1.1. Behavioral results

To contrast the performance between the experimental groups, we first computed the points they obtained over trials (Figure 2A). During the first 200 trials, the Mapping group performed overall better (t(47.98) = 8.06, p < 0.001). This result is expected as the target locations for the Mapping group were only one step away. However, we did not find any significant differences in the overall performance between the groups after trial 200 (t(41.56) = 0.37, p = 0.71). These results could suggest that learning the mapping first did not provide any advantage in the task when the targets were three steps away compared to the Full group. It is important to note, however, that the comparison is not balanced. During the first 200 trials, the Mapping group could generate a maximum of 200 key presses as the trials were terminated with the first key press. In contrast, participants in the Full group did not have this constraint. Indeed, they had a median of 7 key presses per trial during the first 200 trials, and therefore built considerably more experience than the Mapping group. In order to account for this, we grouped the points by key presses rather than by trial (see *Materials and Methods* for details; Figure 2B).

**Figure 2:**
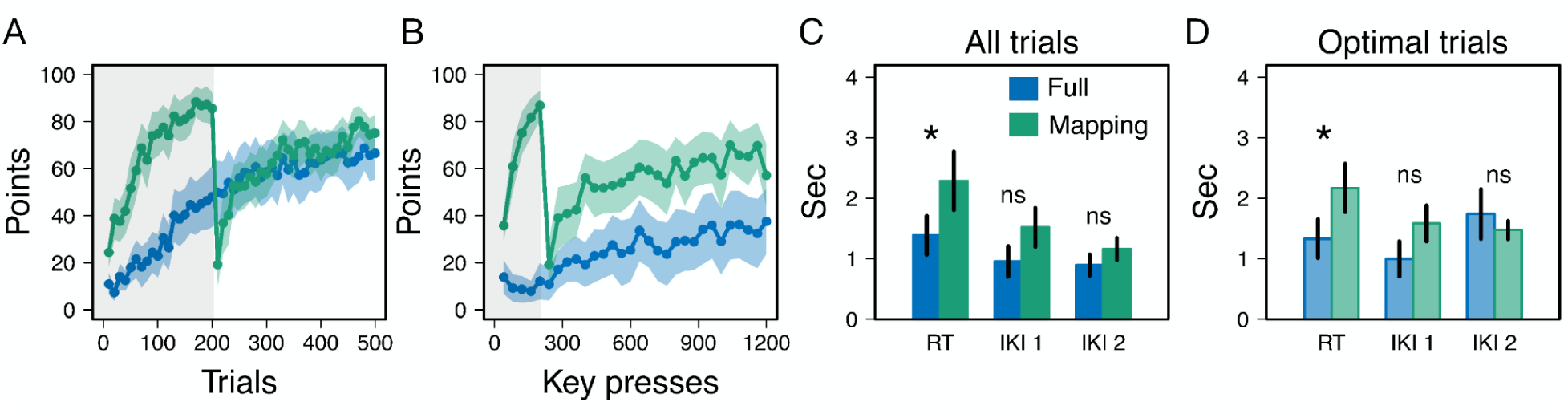
Behavioral results of Experiment 1. **(A)** Points over trials for both experimental groups. The shaded area indicates the first 200 trials (or key presses) where the targets were one step away for the Mapping group. **(B)** Points grouped by key presses. **(C)** RTs and IKIs across all trials. **(D)** Same as in (C) but for optimal trials.

Similar to the comparison across trials, we found that the Mapping group performed better than the Full group for the first 200 key presses (t(35.42) = -14.75, p < 0.001). After key press 200, the performance of the Mapping group significantly dropped (t(39.57) = 11.57, p < 0.001) to the level of the Full group (first bin after key press 200; t(41.87) = -1.39, p = 0.16)).

However participants quickly began to recover their performance, and were significantly better than the Full group in all the following bins (p < 0.001) as revealed by a mixed-effects model analysis. Therefore, when accounting for the number of key presses made in the task, the Mapping group performed remarkably better than the Full group with targets three steps away, suggesting an advantage from learning the mapping first.

To gain insights about the planning processes in our task, we analyzed the reaction time (RT) and inter-key intervals (IKI) when both experimental groups performed the task with the same complexity, i.e., when target locations were three steps away. For the Full group we analyzed the first 300 trials of the task, and for the Mapping group the 300 trials following the exposure to the targets one step away. We performed a 2X3 mixed-design ANOVA with the experimental group as between-subjects factor (Full and Mapping) and time interval as within-subjects factor (RT, first IKI 1 and second IKI). We found that the Mapping group took, in general, took longer to generate their responses as revealed by a significant main effect of the experimental group (F(1,48) = 9.75, p = 0.003, η^2^_*p*_ = 0.17; Figure 2B). Pairwise comparison with Bonferroni correction revealed that RTs were significantly higher in the Mapping group (p = 0.03) but we found no differences between the groups in the first (p = 0.09) or second (p = 0.5) IKI.

We conducted the same analysis for both RT and IKI, focusing exclusively on trials where participants reached the target optimally, i.e., using three moves. These trials were most representative of planning given that the probability of arriving at the target this way without already knowing the mapping is considerably low (see Table S1). Similar to the results for all trials, we found a significant main effect of the experimental group (F(1,48) = 7.85, p = 0.007, η^2^_*p*_ = 0.14; Figure 2C), where the Mapping group generally had higher RTs. Following the same pairwise comparison method, we found that RTs were significantly higher in the Mapping group (p = 0.02) but there were no differences in the first (p = 0.08) or second IKI (p = 0.99). In addition, this time we found that the second IKI in the Full group — which corresponds to the response prior to reaching the target — was significantly higher than the first IKI (p = 0.03). We did not find this difference for the Mapping group (p = 0.99). The latter result indicates that participants in the Full group potentially paused to re-plan the last move before reaching the target.

In summary, our behavioral results show that learning the mapping of the task first, can bring significant performance benefits in subsequent decisions involving planning. In addition, the response time analyses showed that participants in the Mapping group took significantly longer to initiate their responses, while the Full group paused before generating the last response for the trials most representative of planning. This last finding suggests that, while the Mapping group may have planned the whole action sequence beforehand, the Full group most likely relied more on intermediate feedback and replanned before the last move.

#### 3.1.2. Modeling results

We evaluated seven computational models with the goal of explaining the differences in performance between the experimental groups (See *Materials and Methods* for details). These models were inspired by previous research demonstrating the influence of three main variables on decision making: planning (Hunt et al., 2021; Mattar and Lengyel, 2022; Tolman, 1948), reward learning (Sutton and Barto, 1998; Schultz et al., 1997), and action perseveration (Miller et al., 2019, 2021; Daw et al., 2011; Thorndike, 1911). We represented planning using a model-based (MB) algorithm which implements A* search (Hart et al., 1968), and learns the motor mapping between the keys and the state transitions using state prediction errors (Lee et al., 2014). For reward learning (MF), we implemented SARSA (Sutton and Barto, 1998), a widely used temporal difference model which updates state-action values based on the points obtained in the task. For action perseveration, we implemented a model based on Miller et al. (2021) where the value of actions depends on how frequent they have been repeated in the past. In addition, we consider four hybrid models represented as linear combinations of the aforementioned models.

Given that the MB algorithm captures our main variables of interest (the mapping learning and planning), we predicted that the differences in performance could manifest in elements of this model. In particular, we hypothesized that, given the faster improvement in the task by the Mapping group, they would show higher learning rates about the state transitions compared to the Full group.

We followed the method from Stephan et. al (2009) to obtain the probability that a subject selected at random from our experimental groups, was best described by the proposed models. This probability was 99% and 85% in the Full and Mapping groups, respectively, for a two-component hybrid model incorporating model-based planning and action perseveration (MB-Persev; Figure 3A). This model was able to capture on average 47% (Full group) and 57% (Mapping group) of the explainable variability in our data as compared to an information-theoretic near upper bound (Shen and Ma, 2016; Grassberger, 1988, 2003), where 0% represents chance level.

**Figure 3:**
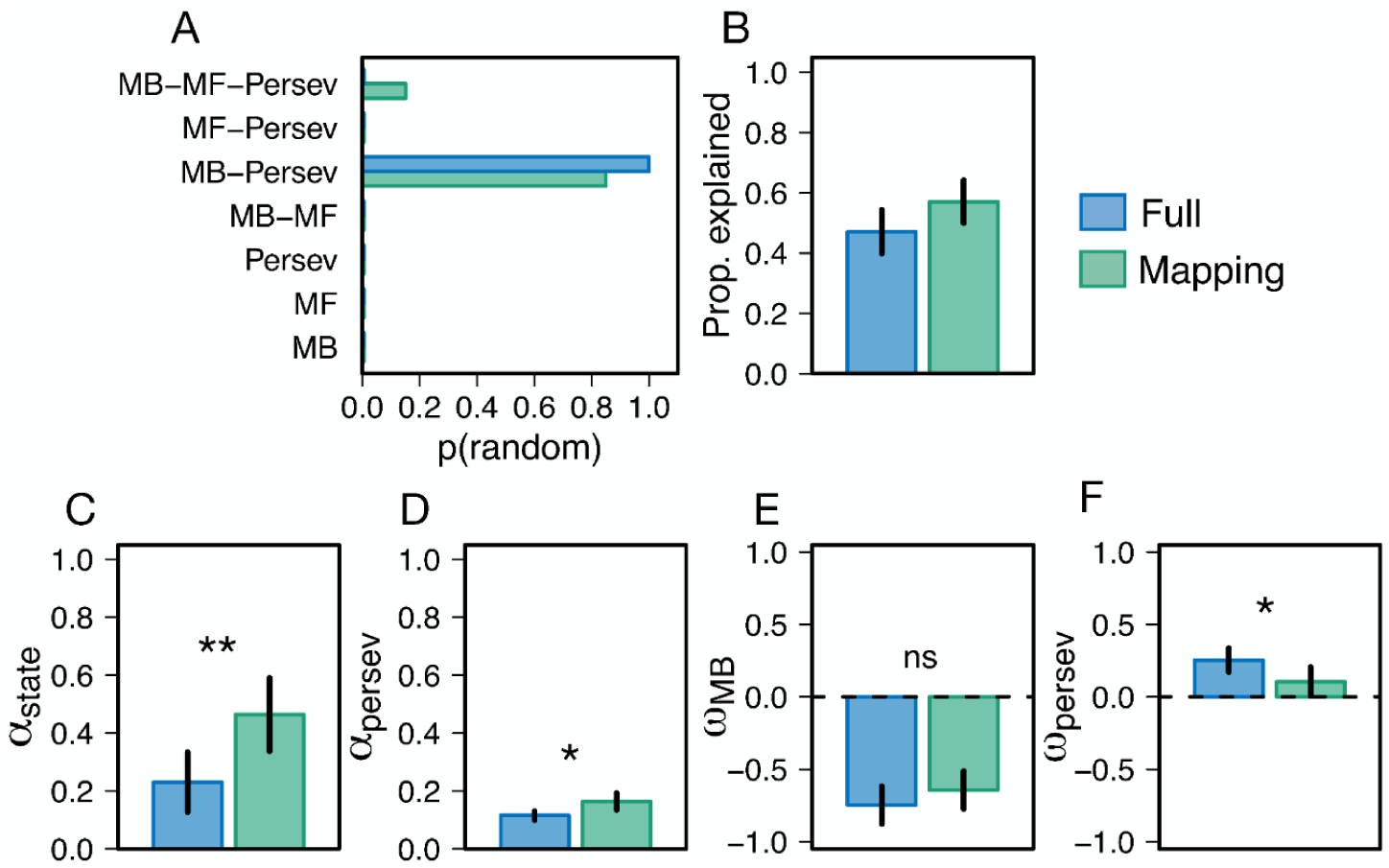
Model results of Experiment 1. **(A)** Probability of obtaining each of the models for a randomly selected participant in the Full and Mapping groups. **(B)** Proportion of the variability in the data explained by the best-fitting model (MB-Persev). **(C-F)** Individual parameter values in the MB-Persev for each of the experimental groups (α_*state*_ = learning rate for the state transitions; α_*persev*_ = learning rate for previous responses; ω_*Persev*_ = weight of the perseveration component; ω_*MB*_ = weight of the model-based component).

As predicted, we found that the Mapping group had significantly higher learning rates for the state transitions, i.e. the motor mapping (α_*state*_ parameter; t(46.2) = -2.94, p = 0.005). In addition, we found that responses that were more distant in the past, had a greater influence for the Full group as indicated by significantly lower values of the parameter α_*persev*_ (t(36.36) = -3.02, p = 0.004). Similarly, we found that the weight of the perseveration component in the hybrid model, ω_*persev*_, was significantly higher for the Full group (t(46.37) = -2.35, p = 0.02). We did not find any significant differences in weight for the model-based component, ω_*MB*_, where its negative values indicate preference for options that incur overall less cost (see *Materials and Methods* for details). Briefly, the cost of an action is determined by the A* search algorithm, which estimates the cost of reaching the target if that action is taken, and this cost is adjusted based on the uncertainty of the mapping.

Overall, our modeling results indicate that better performance of the Mapping group when encountering targets that required sequential decisions, could be attributed to multiple factors. First, as shown by differences in the learning rates for state transitions, the initial training without planning in the Mapping group, could have allowed them to learn the mapping between the keys and the state transitions faster. This, in turn, would have facilitated planning in subsequent stages of the task. Furthermore, the Mapping group had a weaker influence from past responses (Figure 3D) and were less perseverative (Figure 3F) compared to the Full group. Therefore, it is likely that the Mapping group relied more heavily on planning processes rather than habitual responses, which is as also suggested by their higher RT (Figure 2C).

While the sequential nature of our task suggests that planning plays an important role for successful performance, Experiment 1 does not directly manipulate task variables to measure planning. In Experiment 2, we sought to investigate this more directly by varying the distance to the target (i.e., the planning horizon), which previous work has shown to affect the response time associated with the increased demands of planning (Jensen et al., 2024).

### 3.2. Experiment 2

In this experiment, we directly assessed planning in our grid navigation task by manipulating the target distance from the cursor. We hypothesized that key signatures of planning, such as increased RT with the target distance (Jensen et al., 2024) would manifest, and that participants’ performance would decrease for targets further apart due limitations in planning depth.

As in Experiment 1, participants performed 500 trials of the task. However, this time the target could appear from one to four steps away from the start location (Figure 1B). In addition, we incorporated “planning blocks” every twenty trials, where movement and performance feedback was omitted (see *Materials and Methods* for details). The aim of these blocks was to prevent participants from relying on online path corrections, encouraging planning of the whole sequence prior to initiating responses. We hypothesized that this manipulation would maximize the effects generated by the different planning horizons. We refer to the trials in the planning block as “no-feedback trials” and to the trials with feedback as “feedback trials”.

#### 3.2.1. Behavioral results

In order to assess participants’ performance in the task, we used a 4X2 repeated measures ANOVA, where we found a significant main effect of target distance (one to four steps away; F(3,72) = 60.28, p < 0.001, η^2^_*p*_ = 0.71) and trial type (feedback and no-feedback; F(1,24) = 61.75, p < 0.001, η^2^_*p*_ = 0.72) in the points obtained, which indicated that participants performed overall better in the feedback trials and at shorter target distances (Figure 4A). Post hoc pairwise comparisons (Bonferroni corrected), revealed that performance was better in the feedback than in the no-feedback trials for targets at two (p = 0.002), three (p < 0.001) and four (p < 0.001) steps away, but not for targets at one step away (p = 0.85). In addition, we found that, for both types of trials, performance decreased significantly from a given target distance to the subsequent target distance but only up to three steps away (feedback trials: p_1-2_ < 0.001, p_2-3_ = 0.02; no-feedback trials: p_1-2_ < 0.001, p_2-3_ < 0.001). No significant difference in performance was found between targets that were three and four steps away (feedback trials: p_3-4_ = 0.99; no-feedback trials: p_3-4_ = 0.99).

**Figure 4:**
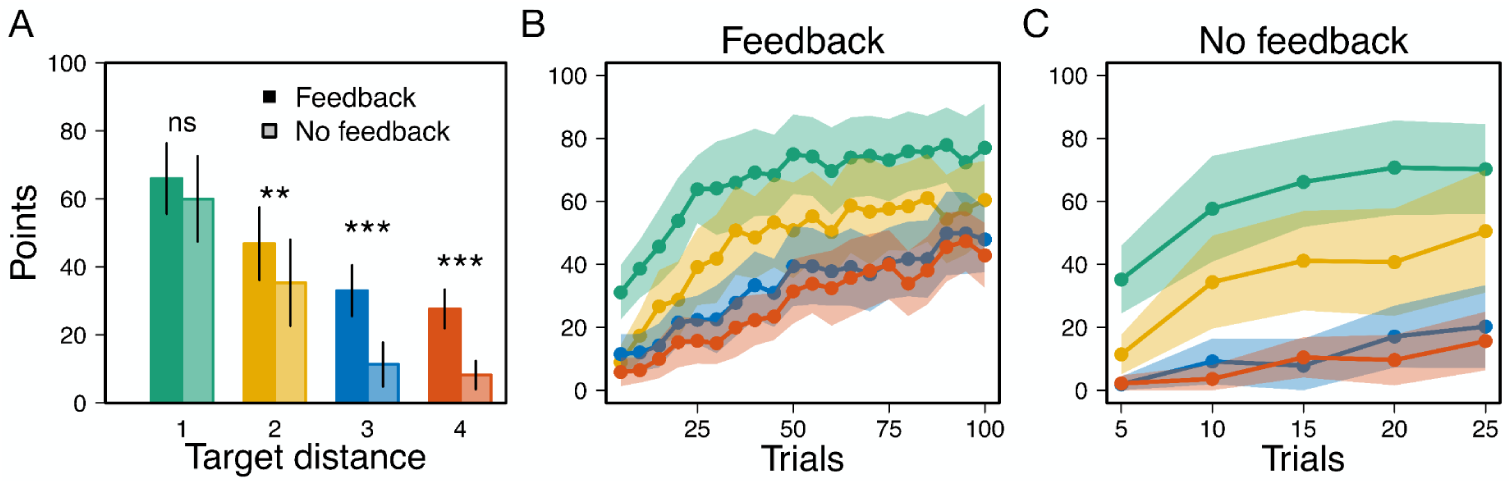
Participants performance in Experiment 2 for the different planning horizons. **(A)** Points across the experiment in the feedback and no-feedback trials. **(B)** Points over trials for the feedback trials. **(C)** Points over trials for the no-feedback trials.

We also assessed whether performance improved across the experiment for the different planning horizons. To do so, we implemented a linear mixed-effects model which included a fixed effect for trial number and a random intercept for subjects. The results revealed a significant effect of trial number on the points obtained for both the feedback (p < 0.001, for all horizons; Figure 4B) and no-feedback trials (p_one_ = 0.009; p_two_ = 0.01; p_three_ = 0.04; p_four_ = 0.01; Figure 4C) which indicates that the performance improved over the course of the experiment for all planning horizons. While performances improved with training, it continued to be better for lower planning horizons.

Importantly for this experiment, we also examined response times (RT and IKI) for the different planning horizons (Figure 5). First, we compared RT for the feedback and no-feedback trials. We carried out a 4X2 repeated measures ANOVA where we found a significant main effect of target distance (F(3,72) = 8.35, p < 0.001, η^2^_*p*_ = 0.25) and trial type (F(1,24) = 15.16, p < 0.001, η^2^_*p*_ = 0.38) indicating that RTs were overall higher for the no-feedback trials (Figure 5B) and differed by target distance. Post hoc pairwise comparisons revealed that RTs were significantly higher in the no-feedback trials for targets at two (p = 0.04), three (p = 0.003) and four steps away (p = 0.004), but not for targets one step away (p = 0.99), which corroborates the increased complexity of multi-step planning without online information.

**Figure 5:**
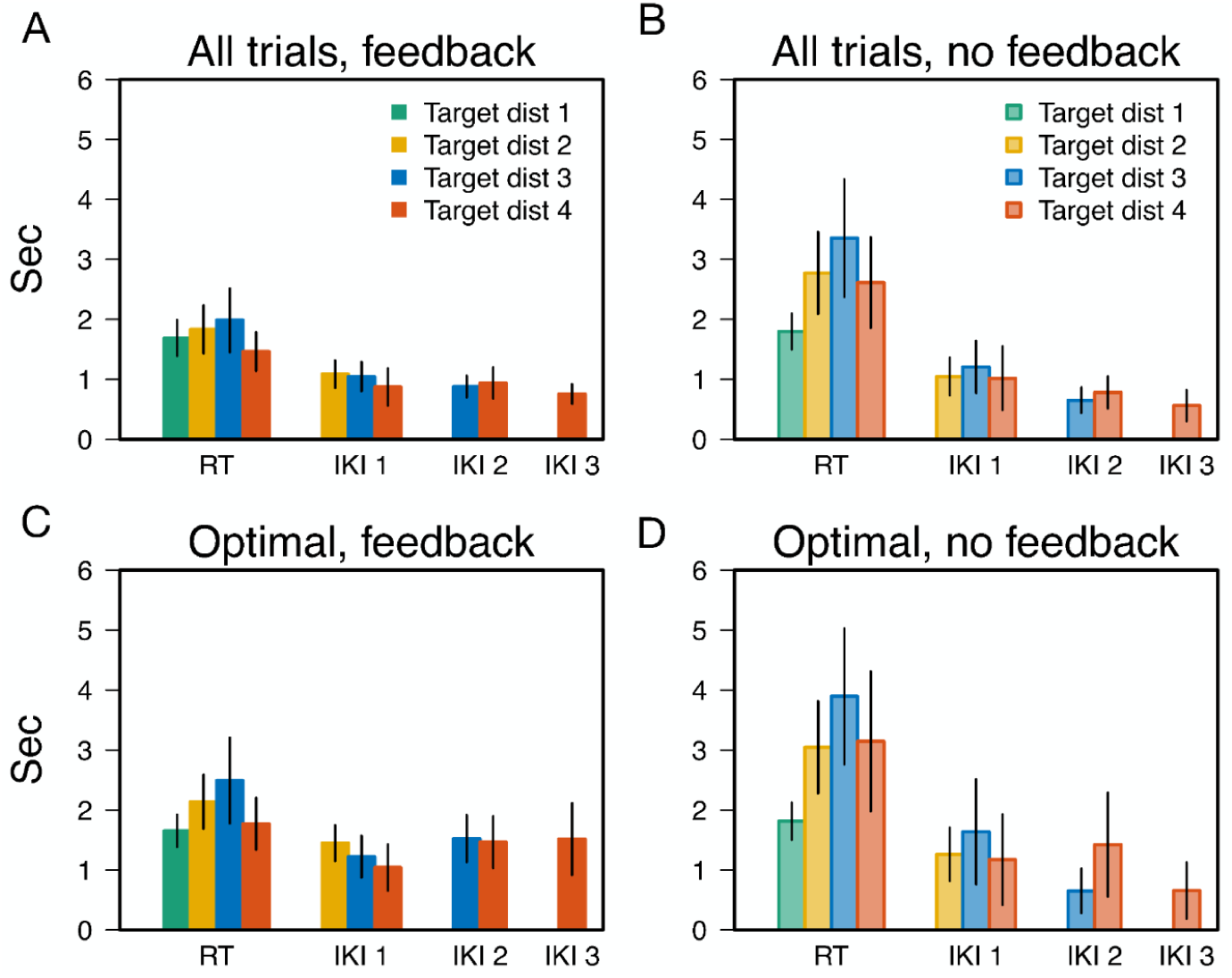
Response times in Experiment 2 for the distinct planning horizons. **(A)** RTs and inter-key intervals for all feedback trials. **(B)** Same as in (A) but for no-feedback trials. **(C)** RTs and inter-key intervals for optimal arrivals in feedback trials. **(D)** Same as in (C) but for no-feedback trials.

Additionally, in the trials with no-feedback, where we expected planning to manifest more strongly, we found a significant positive correlation between RT and target distance up to targets three steps away (t(73) = 3.2, r = 0.35, p = 0.001) but not up to targets four steps away (t(98) = 1.89, r = 0.18, p = 0.06). This difference may indicate a potential planning limit in our task, which we investigate further in our modeling analysis (see below). For the feedback trials, we found no significant correlation between RT and target distance, whether up to three steps away (t(73) = 1.03, r = 0.12, p = 0.3) or four steps away (t(98) = -0.59, r = -0.06, p = 0.55), suggesting that they may have been re-planning later on in the sequence and may be the explanation for the lack of difference in performance between distances 3 and 4 steps away.

We also examined response times for optimal trials, where we assumed participants already knew the mapping and mostly relied on planning (Figure 5C and 5D). Importantly, our sample size was smaller given that not all subjects managed to arrive at the target optimally, particularly for no-feedback trials. Similar to the analysis for all trials, we found a significant main effect of target distance (F(3,59) = 10.49, p < 0.001, η^2^_*p*_ = 0.34) and trial type (F(1,18) = 8.36, p = 0.009, η^2^_*p*_ = 0.31). Notably, post hoc pairwise comparisons revealed that RTs were only marginally higher in the no-feedback trials for the targets at three steps away (p = 0.52), but not for the rest of the distances. This suggests that planning was similar between feedback and no-feedback trials when considering optimal performance. We also found that, for the optimal feedback trials, RT increased significantly up to targets three steps away (t(73) = 2.52, r = 0.29, p = 0.01; Figure 5C), but not up to targets four steps away (t(98) = 0.92, r = 0.09, p = 0.35). For the optimal no-feedback trials, RT significantly increased up to three (t(73) = 4.74, r = 0.54, p < 0.001) and four steps away (t(98) = 3.67, r = 0.42, p < 0.001; Figure 5D).

We did not find any significant difference in the IKI between the trial types or target distances. However, we did find that overall, RTs were higher than the IKI for both feedback (t(38.44) = 4.06, p < 0.001) and no-feedback trials (t(32.28) = 5.42, p < 0.001). In addition, the difference between RT and IKI was significantly larger for the no-feedback trials (t(33.74) = -4.12, p < 0.001), which likely reflected their greater involvement in planning prior to generating the action sequence.

In summary, our behavioral results showed that participants improved performance over the course of the experiment for all planning horizons in both feedback and no-feedback trials. However, their performance decreased for more distant targets and plateaued at targets three steps away. In addition, the fewer points and higher RT in the no-feedback trials for target distances greater than one, highlighted the increased complexity of performing multi-step planning prior to action execution, compared to using online feedback. Lastly, the increase of RT up to targets three steps away in the no-feedback trials, suggested that participants might plan up to this limit. We explore this idea further in our computational modeling analysis.

#### 3.2.2. Modeling results

Based on the modeling results of Experiment 1, where the MB-Persev model best described the data, we considered variations of that model which differed in their planning depth (See *Materials and Methods* for details). Specifically, we evaluated models that estimate the value of actions based on simulations ranging from one to four steps ahead. Figure 6A shows an example of how the performance of the model can differ depending on the planning depth. If the model considers only one step ahead, it may select suboptimal actions that appear to reduce the distance to the target (green node) based on its heuristic search. However, it can overlook actions that initially seem to move away from the target but ultimately reach it in subsequent steps (gold trajectory). Such options will only be preferred if a deeper search is performed. This model is able to predict the decrease in performance for the distinct planning horizons similar to what we observed in participants (Figure 5B). While performance generally decreases with the distance to the target, this effect is attenuated as the depth of the algorithm increases. These simulations were generated using the mean of the best fitting parameters from the Full group in Experiment 1.

**Figure 6:**
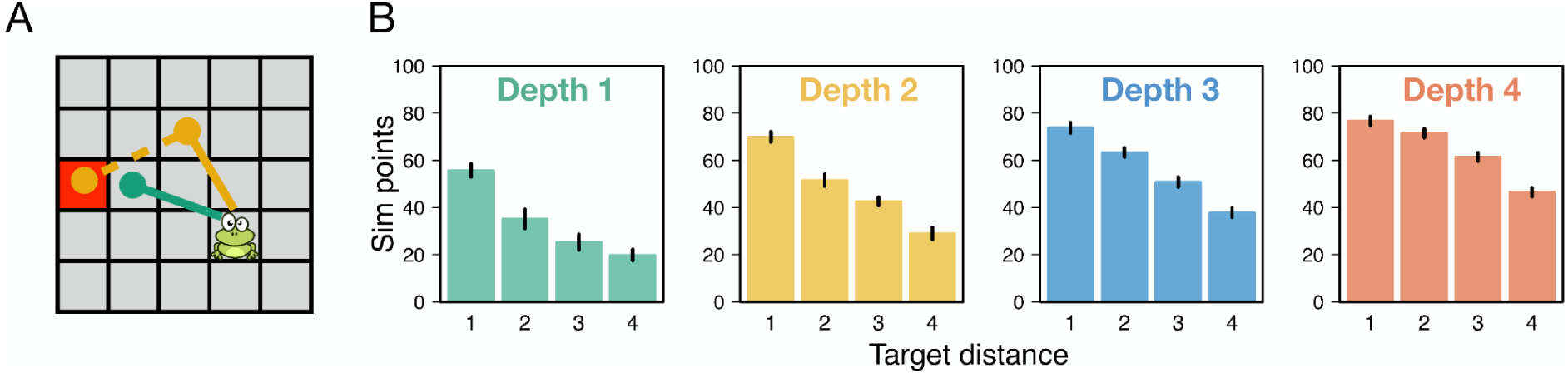
Simulations of the MB-Persev model with different depths of search. **(A)** Example of the preferred moves from an algorithm with depth 1 (green trajectory) or depth 2 (gold trajectory). While an algorithm with depth 1 prefers the option that minimizes the immediate chessboard distance to the target, an algorithm with depth 2 prefers an option which is not as close to the target immediately, but leads to it in the next move. **(B)** Simulated points for the different target distances for different depths of search. Overall, performance decreases for more distant targets, but this effect is attenuated as the depth of search increases.

As in Experiment 1, we computed the probability that a random participant from the population was best described by each of our models (Stephan et al., 2009). We found that this probability is 60% for the MB-Persev model with depth 3, followed by 20% with depth 2, and 18% with depth 4. In addition, we found that the fest fitting model (MB-Persev depth 3) was able to capture a mean of 40% of the explainable variability in the feedback trials (Shen and Ma, 2016; Grassberger, 2003; Figure 7B).

**Figure 7:**
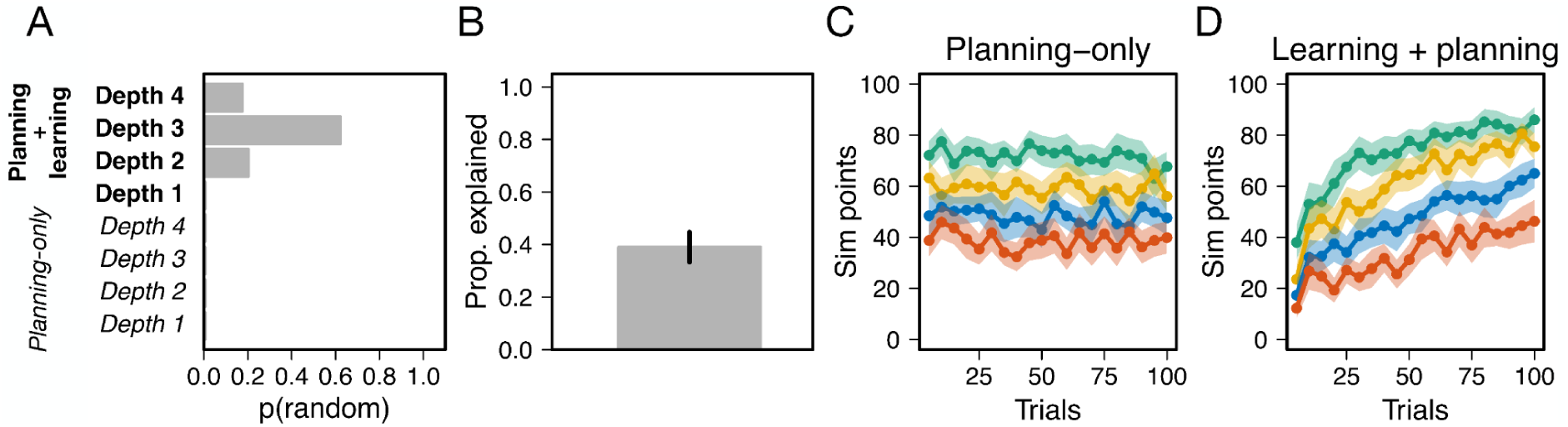
Model results of Experiment 2. **(A)** Probability of obtaining each of the MB-Persev models with different depths for a randomly selected subject. **(B)** Proportion of the variability explained by the MB-Persev model with depth 3. **(C)** Simulated points over trials for the Planning-only model (MB) with depth 3 (green = horizon 1; gold = horizon 2; blue = horizon 3; orange = horizon 4). **(D)** Simulated points for the Learning + planning (MB-Persev) with depth 3.

In order to test the importance of mapping-learning in our task, we also considered four “planning-only” models which do not incorporate any learning about state transitions or previous responses into their computations. In such models, the value of actions is entirely the result of the tree search from the planning algorithm with varying depths.

We fitted all models using the responses from the feedback trials and used the parameter estimates to describe the performance in the no-feedback trials. While these planning-only models are able to capture differences in performance for the distinct planning horizon, it assumes that performance remains stable across the experiment (Figure 7C) and does not reflect the learning time course of performance of the participants (Figure 4B and 4C). In contrast, the MB-Persev depth 3 model, which incorporates learning about the state transitions and about previous actions, is able to qualitatively capture the improvement in performance across the experiment (Figure 7D).

We further assessed the performance of the MB-Persev depth 3 model in the no-feedback trials by simulating response probabilities using the observations that participants had in those trials and best-fitting parameters from the feedback trials (Figure 8). We only considered responses up to the planning depth for the given trial. For example, if the target was two steps away, we only considered the likelihood of the first two responses. With this, we aimed to exclude responses resulting from guessing, which were likely to happen as the number of responses increased due to the uncertainty about the cursor location. Overall, we found that the model was reasonably accurate at describing responses for targets one step away.

**Figure 8:**
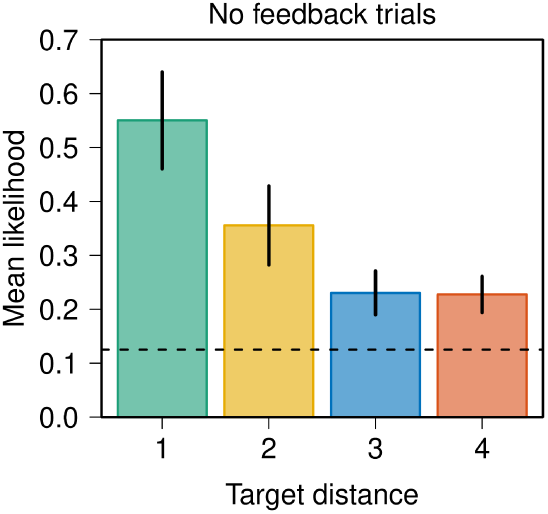
Model predictions for the no-feedback trials in Experiment 2. The mean likelihood for the no-feedback trials was obtained using the best-fitting parameters of the model and as many responses as the target distance for the given trial.

However, its accuracy decreases for longer distances, although it performed greater than chance in all cases (p < 0.001; Figure 8)

In summary, the modeling results of Experiment 2 support our findings in performance and RT, suggesting that participants may plan between 2 to 4 steps ahead in our task, with the majority of participants performing 3-step planning. In addition, we found that a model that incorporates learning and planning into its computations, best captures the performance improvement over trials as well as the decrease in performance for longer planning horizons.

## 4. Discussion

When acquiring novel motor skills, humans face the double challenge of learning what the outcomes of their actions are and how to use them sequentially to achieve future goals. The study of this matter lies at the heart of motor learning and planning research. However, both fields have addressed the topic mostly from separate fronts. Much of motor learning research has focused on sequence learning tasks where the planning component is constrained or limited by the experimenter. On the other hand, planning studies rarely focus on situations where the state transitions require motor learning. In two experimental studies, we aimed to bridge this gap by using a grid navigation task where participants have to learn a novel motor mapping based on the Knight piece from chess, and use it sequentially to arrive at target locations.

In Experiment 1, we showed that performance in this task can benefit from learning the motor mapping first, in the absence of planning. As revealed by our best-fitting model, this training regime can lead to increased learning rates about the state transitions, as well as to a lower influence from habitual responses. The increased performance in a task by breaking it down into its components has been reported previously in the so-called part-whole training (Mané and Donchin, 1989; Park et al., 2010). Practicing a complex component of a perceptual-motor task, such as learning the motor mapping, can yield benefits when the task is later performed as a whole, which can be a useful strategy for acquiring complex motor skills, like in sports. Crucially, our findings suggest that this improvement is partly due to increased learning rates in the state transitions associated with the motor mapping. While we explicitly designed the task so the Mapping group learned the motor mapping first, it raises the interesting question of how humans break a task into subcomponents on their own (Correa et al., 2023). Our results suggest, however, that without an explicit design, it may be considerably harder to break down a task as indicated by the significantly lower performance of the Full group.

Additionally, in Experiment 2 we showed that participants’ performance decreased for longer planning horizons; however, performance significantly improved over the course of the experiment in all of them. This pattern was best explained by a model that incorporates both learning the motor mapping and planning with a depth of three steps ahead. Additional support for this planning depth was found in the significant increase in RT for targets up to three steps away in the no-feedback trials, where planning was expected to manifest more strongly. Although similar results have been reported in “planning-only” tasks (Jensen et al., 2024), our results highlight the critical role of motor learning to capture the improvement of participants across the experiment. In a previous study we have addressed whether prior experience in a planning-only version of our task could benefit the performance when participants learned the mapping and planned, for example, by reducing the cost of planning. However, it did not confer any benefit (Velazquez-Vargas and Taylor, 2023).

Based on previous work on decision-making, we believe the main results from Experiment 1 and 2 could be in part attributed to the distinct working memory loads in the task (Collins and Frank, 2012; Yoo and Collins, 2022). Indeed, in the seminal work from Collins and Frank (2012), the authors show that in an instrumental learning task humans improved their performance at slower rates for larger set sizes of the learned stimuli. A similar effect can be thought to occur in Experiment 1, where the Full group needs to learn an eight-key mapping while potentially attempting to build a decision tree for planning (Huys, 2012). In contrast, the Mapping group only needs to learn the mapping during the first 200 trials, after which they can focus exclusively on planning. Evidence that the Mapping group had a greater involvement in planning when the targets were three steps away, comes from their higher RT compared to the Full group (Figure 2B and 2C).

In addition, the differences in performance for the distinct planning horizons in Experiment 2 (Figure 4B) are remarkably similar to the patterns found by Collins and Frank (2012) for the different set sizes of stimuli. These results suggest that working memory and planning are closely related processes, an idea that has been addressed in recent work (Bottcher et al., 2021; Zhuojun et al. 2023). Indeed, It is conceivable that planning, which is typically thought of as building a decision tree (van Opheusden, 2023; Mattar and Lengyel, 2022) is constrained by working memory capacity. For example, it can limit the size or depth of the tree, or the precision of the reward stored at different levels (Zhuojun et al. 2024). Such constraints can underlie the use of heuristics to reduce the computational demands of planning such as pruning (Huys et al., 2012, 2015) or truncation (Keramati et al, 2016; Krusche et al., 2018) of the decision tree. In our task, our planning model (A*) was constrained in two ways that may reflect the limited cognitive resources in humans. First, we incorporated a limited planning depth. Second, through the incorporation of the chessboard distance as a heuristic function in A*, we specified that planning is more likely to occur in the direction of the goal, rather than in any direction such as in uninformed algorithms like Breadth-First Search. These modifications can considerably reduce the working memory load.

While planning depth varies by task, our estimate of three steps ahead lies at the lower end of previously reported planning depths (Huys et al., 2015; van Opheusden, 2023; Arad and Rubinstein, 2012; Snider et al., 2015; Ariani et al., 2021). This can reflect the complexity of planning using the Knight rule, which incorporates non-linear transitions between states and may increase working memory load. Future work can explore how planning differs for mappings with distinct geometries in the state-transitions. For example, by comparing the planning depth while using the Knight mapping and the King mapping. Although both mappings have the same number of actions (eight moves), the latter moves to the adjacent states, which can reduce the complexity of planning and increase the planning depth.

It is important to note that while our best-fitting model qualitatively captures the behavioral patterns in our task, it still falls short in describing the full extent of our data (Figure 3B and Figure 7B). This could be in part due to cognitive processes that were not specified in our models. However, it could also be attributed to guessing responses, for example, due to attentional lapses. While we did not explore the extent to which participants guessed in our tasks, it could be addressed by extending the model to incorporate a contaminant process, which is common in working memory experiments (Adam et al., 2017; Velàzquez-Vargas and Taylor, 2024). Additionally, other planning algorithms could be considered such as Monte Carlo Tree Search, a common planning algorithm in board games (Silver et al., 2016, 2017), which has also been implemented to model how humans plan (Élteto et al., 2023; Krusche et al., 2018).

Finally, while our results provide insights into how motor skills are acquired in discrete state spaces, such as in video games, little is known about how this process occurs in continuous domains. Crucially, a great variety of sequential motor skills, such as playing a musical instrument or sports, involve the control of continuous actions in continuous spaces. These scenarios likely involve the use of internal forward models to simulate the outcomes of candidate actions in combination with internal models about how the world works (McNamee and Wolpert, 2019). Similar to non-motor planning tasks, the dimensionality of the problem suggests the use of constrained algorithms which, in combination with the use of stereotypical or cached solutions (Jax and Rosenbaum, 2007), can make planning tractable.

## 5. Conclusion

Overall, the present studies show the critical role of integrating motor learning into planning tasks, which is a crucial step to understanding how humans acquire a wide and complex set of motor skills throughout life.

## Data availability statement

Supplementary material of this article, including code and data, is available on the Open Science Framework at https://osf.io/wm5fu/. A preliminary version of this work was presented at the 45 Annual Meeting of the Cognitive Science Society.

## Acknowledgements

We thank the members of the IPA Lab for helpful discussions.

## Supplementary material

**Figure S1:**
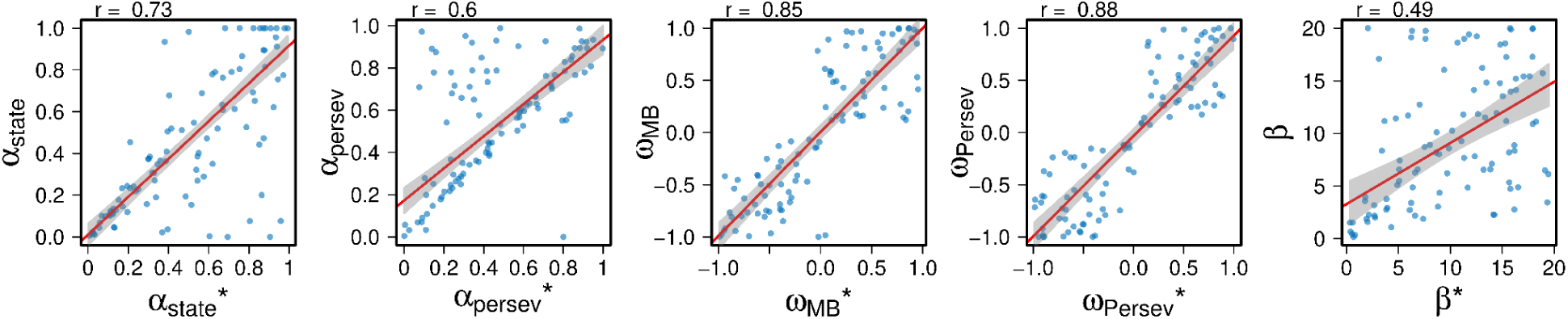
Parameter recovery of the best fitting model (MB-Persev) using a depth of three steps ahead.

**Figure S2:**
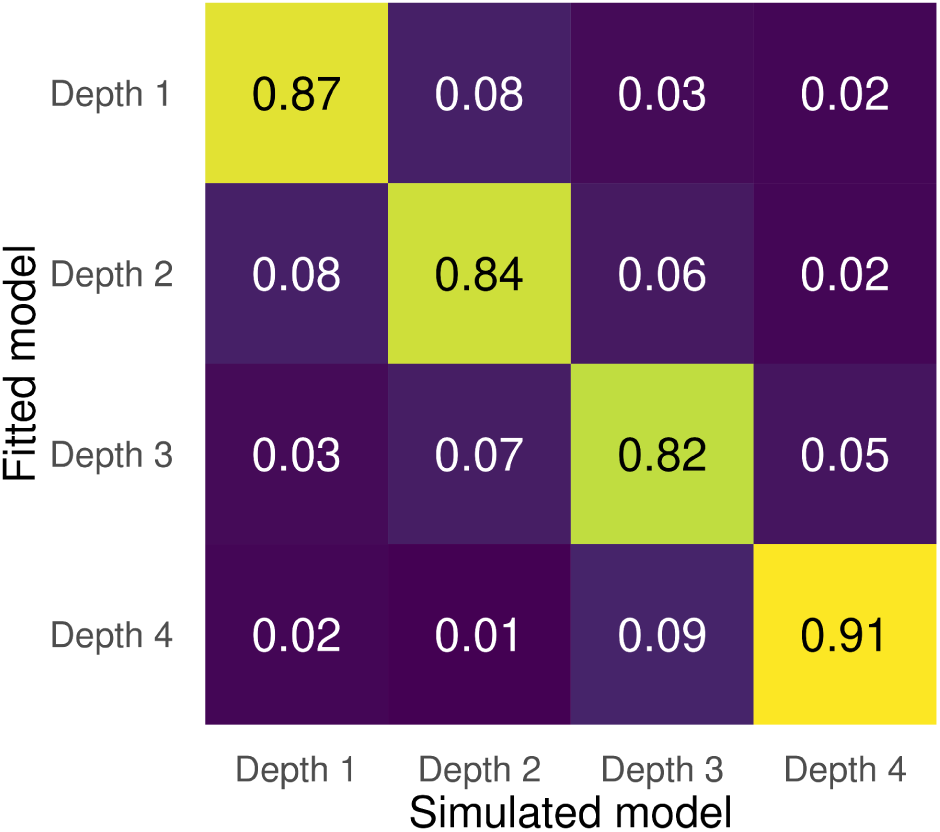
Confusion matrix with model recovery results. Each row and column shows the MB-Persev model with different depths of search. Numbers inside the cells represent the proportion of times that the model in the Y axis best recovered the data generated by the model on the X axis according to BIC.

**Table S1.**
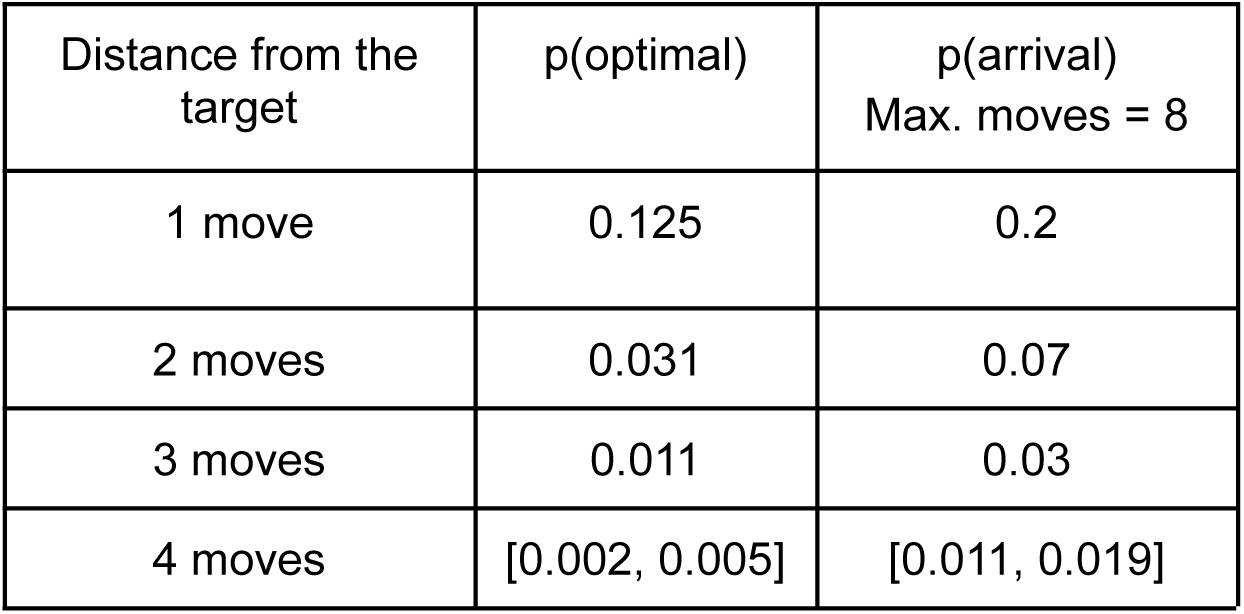
Probability of randomly reaching the target with the optimal number of moves (second column) and within eight moves (third column). These probabilities were computed using Breadth First Search (BFS). A maximum of eight moves was considered based on the observation that participants on average made 7.43 moves across both experiments

| Distance from the target | p(optimal) | p(arrival)<br>Max. moves = 8 |
| --- | --- | --- |
| 1 move | 0.125 | 0.2 |
| 2 moves | 0.031 | 0.07 |
| 3 moves | 0.011 | 0.03 |
| 4 moves | [0.002, 0.005] | [0.011, 0.019] |

